# Exploring the Causal Effects of Shear Stress Associated DNA Methylation on Cardiovascular Risk

**DOI:** 10.1101/2020.08.07.241554

**Authors:** Ruben Methorst, Gert Jan de Borst, Gerard Pasterkamp, Sander W. van der Laan

## Abstract

**Background and aims:** Atherosclerosis is a lipid-driven inflammatory disease presumably initiated by endothelial activation. Low vascular shear stress is known for its ability to activate endothelial cells. Differential DNA methylation (DNAm) is a relatively unexplored player in atherosclerotic disease development and endothelial dysfunction. Literature search revealed that expression of 11 genes have been found to be associated with differential DNAm due to low shear stress in endothelial cells. We hypothesized a causal relationship between DNAm of shear stress associated genes in human carotid plaque and increased risk of cardiovascular disease.

**Methods:** Using Mendelian randomisation (MR) analysis, we explored the potential causal role of DNAm of shear stress associated genes on cardiovascular disease risk. We used genetic and DNAm data of 442 carotid endarterectomy derived advanced plaques from the Athero-Express Biobank Study for quantitative trait loci (QTL) discovery and performed MR analysis using these QTLs and GWAS summary statistics of coronary artery disease (CAD) and ischemic stroke (IS).

**Results:** We discovered 9 methylation QTLs in plaque for differentially methylated shear stress associated genes. We found no significant effect of shear stress gene promotor methylation and increased risk of CAD and IS.

**Conclusions:** Differential methylation of shear stress associated genes in advanced atherosclerotic plaques in unlikely to increase cardiovascular risk.

**Highlights:**

- Plaque-derived DNA methylation in shear stress associated genes shows no significant effect on cardiovascular disease
- Genetic variants in shear stress associated genes affect DNA methylation in human carotid plaque
- Human validation of atherosclerotic associated genes in murine models

## Introduction

Atherosclerosis is a lipid-driven inflammatory disease underlying many cardiovascular diseases, such as coronary artery disease (CAD) and ischemic stroke (IS). Low shear stress is likewise a key player in atherosclerosis and results in endothelial activation, ultimately leading to the initiation and progression of atherosclerotic plaque formation [1,2]. In mice differential DNA methylation (DNAm) at the promoter region of 11 shear stress associated genes (HOXA5, TMEM184B, ADAMTSL5, KLF4, KLF3, CMKLR1, PKP4, ACVRL1, DOK4, SPRY2 [3], and ENOSF1[4]), was shown to alter gene expression and influence endothelial dysfunction [3,5].

However, it is unclear to what extent this applies to humans. It is well established that DNAm regulates gene transcription by modulating the interaction between DNA and chromatin binding proteins [14]. Given that common cardiovascular risk factors, such as smoking [6] and obesity [7–9], are known to associate with DNAm, these risk factors could give rise to aberrant DNAm, thereby impeding physiological regulation of gene expression and negatively impacting atherosclerotic progression. Here, we assess if shear stress could also play a similar role by using stated murine genes using an in silico approach to determine causality between shear stress associated DNAm and cardiovascular risk (Fig. 1). Of course, observed differential effects in shear stress could also be due to reverse causality or residual confounding and genes identified in mouse models might not reflect the human representation of genes affected by shear stress.

**Figure 1:**
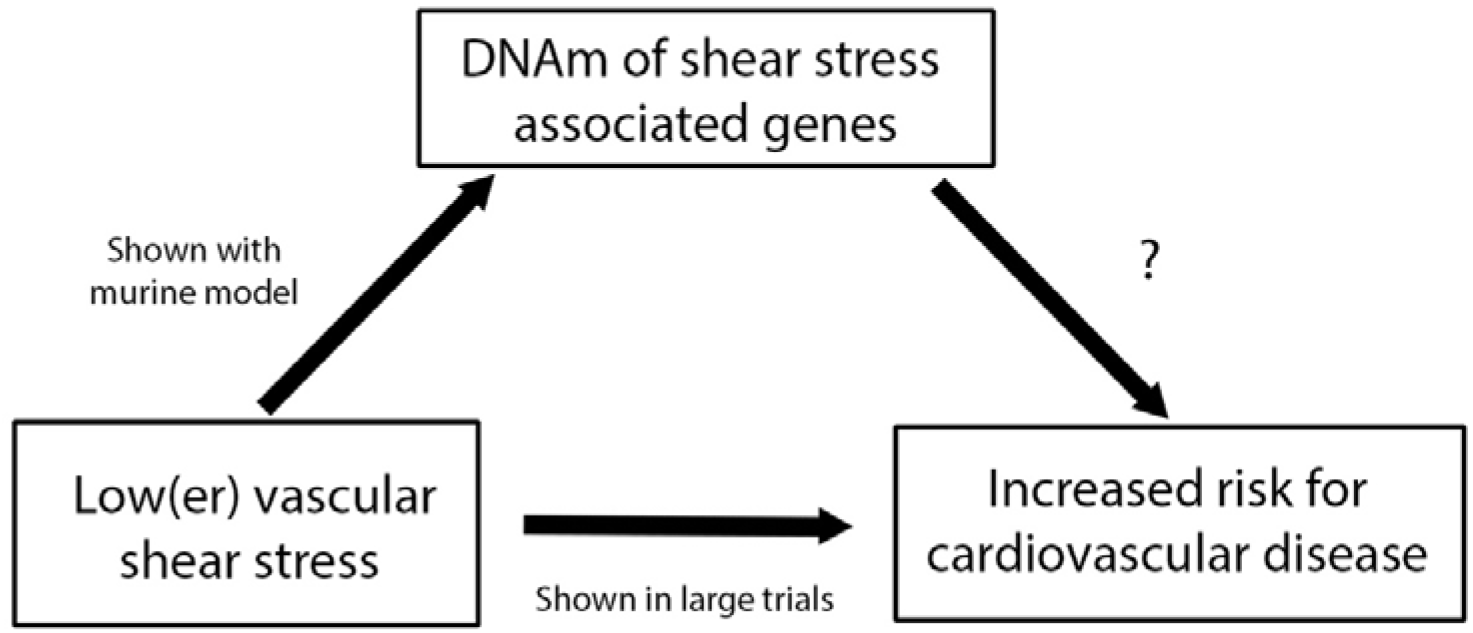
Causal inference scheme of DNAm of shear stress associated genes on cardiovascular risk. It has been shown that a low(er) vascular shear stress is associated with an increased risk for cardiovascular disease in multiple large trials. Dunn et al. showed that a low shear stress results in differential methylation of 11 shear stress associated genes. Here, we explore the final line of causality. The effect of this differential methylation on cardiovascular risk.

To assess the impact of shear stress associated DNAm on cardiovascular disease, we propose to implement Mendelian randomisation (MR) to identify the causal inference between shear stress associated DNAm and cardiovascular outcome. For this, we identified methylation quantitative trait loci (mQTLs) to predict the presence of DNAm using genetic variants, i.e. as input proxy for MR, and calculated causal inference between shear stress associated DNAm and cardiovascular risk.

Much akin randomized clinical trials, MR studies make use of intrinsic properties of the genome for causal inference: as alleles are randomly distributed from parents to offspring at conception, the genetic information is not influenced by disease (reverse causality), or risk factors (residual confounding), and remains largely unchanged throughout life [10,11]. Large-scale genetic analyses of cardiovascular diseases, including CAD [12] and IS [13], and cardiovascular risk factors enables us to infer whether DNAm at shear stress associated genes are causal to such processes, e.g. shear stress results in differential DNAm of certain genes, leading to differential expression adverse for atherosclerotic lesion progression. Determining this causal inference contributes to a better understanding of atherosclerotic initiation, propagation and underlying mechanisms in humans. To this end, we set out to identify genetic variants that predict DNAm, mQTLs, in advanced plaques from the Athero-Express Biobank Study and used these genetic variants to infer causality of DNAm on CAD and IS risk using MR.

## Patients and Methods

### Athero-Express Biobank Study

The Athero-Express Biobank Study (AE, www.atheroexpress.nl) is a longitudinal biobank study including patients that undergo either carotid or femoral endarterectomy in two Dutch tertiary referral centres. The biobank study is ongoing, and its database has been expanding since 2002. A detailed cohort description has been published by Verhoeven et al., 2004 [15]. In this study, genotype, methylation and phenotype data of carotid endarterectomy patients was used. The study was approved by the ethical commission of the participating medical centres. All participants provided informed consent. The study complies with the Declaration of Helsinki.

### DNA isolation

Carotid plaque specimens were removed during surgical intervention and processed following specific guidelines (please refer to Verhoeven et al., 2014). In short, specimens were cut into 5 mm segments and culprit lesions were identified to be fixed in 4% formaldehyde embedded in paraffin. Histological features were scored and remaining segmented were stored at −80 °C until further processing. DNA isolation was performed on these segments according to in-house protocols as described by Van der Laan [16].

### DNA methylation

Isolated DNA samples were randomly distributed on 96-well plates at equalized DNA concentrations of 600 ng. DNA was bisulfite converted using a cycling protocol and the EZ-96 DNA methylation kit (Zymo Research, Orange County, USA). The Infinium HumanMethylation450 Beadchip Array (HM450k, Illumina, San Diego, USA) was used to measure DNA methylation, processing according to manufacturer’s protocol. The HM450K experiment was performed at the Erasmus Medical Center Human Genotyping Facility in Rotterdam, the Netherlands. In total, we collected data from 442 AE patients for the Athero-Express Methylation Study 1 (AEMS450K1) [6].

### Genotyping and imputation

DNA was isolated from stored samples according to the above mentioned protocol and genotyped in two phases (Athero-Express Genomics Study 1 (AEGS1) and Athero-Express Genomics Study 2 (AEGS2)) [16]. Both AEGS1 and AEGS2 samples were genotyped using commercially available genotyping arrays, respectively the Affymetrix Genome-Wide Human SNP Array 5.0 and the Affymetrix Axiom® GW CEU 1 Array. Quality control was performed using community standards and assurance procedures [16,17]. Our reference panel consisted of a merge of phased haplotypes from the 1000 genomes project (phase 3, version 5) [18] and haplotypes from the Genome of the Netherlands (GoNL5) [19] and was imputed using IMPUTE2 [20].

### Methylation quantitative trait loci analysis

We used the QTLToolKit workflow (swvanderlaan.github.io/QTLToolKit/) [21] which leverages QTLtools [22] to identify cis-acting mQTLs in carotid plaques of our genes of interest. The region of interest (ROI) was determined by flanking the outermost DNAm sites (CpGs) of the −2000 transcription start site (TSS) to the first exon by 250 kb upstream and downstream (Suppl. Table 1). We used these ROIs to test for phenotype-genotype pairs, i.e. associations between CpGs and variants. Two passes were performed, an initial pass to get nominal P-values on our dataset and a permutation pass to correct for multiple testing error (FDR < 5%) and get adjusted P-values. We filtered out potential false positives caused by variants affecting the binding of a probe on the array by removing CpG-variant pairs within the same probe and in linkage disequilibrium (LD) with the same probe.

### Two sample Mendelian randomization

To determine causal effect of DNA methylation of shear stress associated genes on CAD and IS we applied the Two Sample Mendelian Randomisation (2SMR) design (using the R-package TwoSampleMR) [10]. The 2SMR design is able to infer causality between an exposure (DNAm) and an outcome (CAD or IS) by using public genome wide association study (GWAS) summary statistics available through the MR-Base platform (http://www.mrbase.org). Variant proxies were used for outcome GWAS variants, if not available in that particular GWAS (LD R^2^ < 0.8). We used GWAS summary-statistics from the CARDIoGRAMplusC4D [12] study for CAD and GWAS summary-statistics from the METASTROKE [13] study for IS. We used the cis-acting mQTLs of plaque tissue as proxy of the exposure (DNAm). Respectively, 3 and 1 Variant(s) passed LD clumping and harmonization to GWAS summary statistics and were used for 2SMR analysis.

### Statistical analysis

Details on the statistical analyses in CARDIoGRAMplusC4D, and METASTROKE were previously described [12,13]. For the discovery of cis-acting mQTLs in carotid plaques, we assumed an additive genetic model and corrected for sex, age, and genotyping array type. To declare a for causal relationship between exposure and the significance was set at p < 0.05. We used Inverse Variant Weighted (IVW) and MR-Egger (intercept) to determine causality. IVW combines ratio estimates of individual genetic variants to a weighted mean, resulting in a consistent estimate of the causal effect, which converges to true values as sample size increases. Therefore, IVW is an efficient analysis method, but it will be biased if only a single genetic variant is invalid. MR-Egger Regression performs a weighted linear regression and if there is no intercept term, it is equal to IVW. A non-zero of the intercept can be interpreted as an estimate of the horizontal pleiotropic effects (an effect not mediated via the exposure) of the genetic variants, indicating directional pleiotropy, and suggesting IVW is biased [23]. Furthermore, MR-Egger can provide a true causal effect if the genetic variant is not independent from the outcome, using the inSIDE (instrument strength independent of direct effect) assumption. mQTL power estimation showed a strong power of 85% and higher at minor allele frequencies (MAF) > 0.06 (Suppl. figure 1).

### Data availability

Scripts available from: https://github.com/rubenmethorst/shear-stress-project. Data available upon request.

## Results

### Common variants predict methylation of shear stress genes

To be able to perform Mendelian randomisation (MR) with DNAm, we identified common genetic variants that are able to predict DNAm in individuals. For this, we genotyped 1,439 individuals from the AE^5^ and extracted DNA from 442 overlapping advanced atherosclerotic carotid plaque samples to assess methylation (Table 1) [6]. We defined regions of interest (ROIs) between the −2,000 transcription start site (TSS) and the first exon for each of the 11 shear stress associated genes (Suppl. Table 1). We used the QTLToolKit [21] and QTLtools^4^ to test for common cis-acting methylation quantitative trait loci (mQTL) within ±250kb of the ROIs and discovered 121,109 potential mQTLs near the 11 genes at nominal p-values (Supplemental excel table). To correct for multiple testing, we performed permutation (adaptively scaled between 1000 and 10,000 permutations) and identified 12 significant cis-mQTLs-CpG pairs at 3 genes (Table 2). Regional association of the highest associated variant-CpG pair corresponding with a shear stress associated gene, shows a strong statistical relationship between rs7235957 and multiple CpG sites in the ENOSF1 promotor (lowest p-value: p= 1.47×10^−38^) (Fig. 2, Table 2). The 12 significant cis-mQTL-CpG pairs are used for MR analyses as genetic instruments for promotor DNAm at shear stress associated genes.

**Table 1.** Baseline characteristics Athero-Express Biobank cohort. Patient characteristics at baseline inclusion. SBP; systolic blood pressure, DBP; diastolic blood pressure, BMI; body-mass index, LLDs; lipid lowering drugs, Ocular; retinal infarction and amaurosis fugax, TIA; transient ischemic attack, and freq; frequency.

| Characteristic | Discovery (n=442) |
| --- | --- |
| Age, y (SE) | 67.9 (9.01) |
| Males (%) | 68.8 |
| SBP, mm Hg (SE) | 156.1 (25.88) |
| DBP, mm Hg (SE) | 82.5 (13.24) |
| BMI, kg/m<sup>2</sup>(SE) | 26.7 (3.94) |
| Smoking (% (freq)) | 40.3 (178) |
| Comorbidities (% (freq)) |  |
| Diabetes Mellitus | 22.6 (100) |
| Hypertension | 87.3 (386) |
| Medication use (% (freq)) |  |
| Hypertensive drugs | 77.4 (342) |
| Anticoagulants | 12.4 (55) |
| LLDs | 3.4 (15) |
| Symptoms (%)<sup>a</sup> |  |
| TIA | 41.4 |
| Stroke | 25.8 |
| Asymptomatic | 14.0 |
| Ocular | 13.1 |
| Other | 5.7 |

<sup>a</sup>symptoms at presentation tertiary referral centre for carotid endarterectomy

**Table 2.** Shear stress related differential DNAm associated p 464 ermutated cis-mQTLs in advanced plaques. Shear stress associated cis-mQTLs in advanced plaques and corresponding CpG sites within the ROIs. MAF > 0.5 was used for analysis. Chr; Chromosome, BP; chromosome location relative to 1000 Genomes Project (Nov 2014, Hg19), CAF; coded allele frequency, HWE; Hardy-Weinberg-Equilibrium, INFO; imputation quality, Gene Name; refSeq (GRCh37/hg19) canonical genes from UCSC, SE; Standard Error, Perm P-value; permutation P-value.

| CpG | CpG Position | Variant | Chr | BP | Other Allele | Coded Allele | CAF | HWE | INFO | Gene Name | Beta | SE | Nominal P-value | Perm P-value |
| --- | --- | --- | --- | --- | --- | --- | --- | --- | --- | --- | --- | --- | --- | --- |
| cg07100532 | TSS1500 | rs7235957 | 18 | 717,229 | T | C | 0.544 | 0.425285 | 0.9793 | *ENOSF1* | 0.794 | 0.054 | 1.12E-48 | 1.47E-38 |
| cg26147554 | TSS200 | rs7235957 | 18 | 717,229 | T | C | 0.544 | 0.425285 | 0.9793 | *ENOSF1* | 0.752 | 0.058 | 2.95E-39 | 9.20E-33 |
| cg16112050 | TSS1500 | rs7235957 | 18 | 717,229 | T | C | 0.544 | 0.425285 | 0.9793 | *ENOSF1* | 0.478 | 0.038 | 6.51E-36 | 6.91E-30 |
| cg15158376 | TSS200 | rs1061035 | 18 | 722,118 | A | G | 0.121 | 0.535303 | 0.9902 | *ENOSF1* | 0.805 | 0.064 | 9.51E-37 | 1.69E-29 |
| cg00955482 | TSS200 | rs2741188 | 18 | 708,299 | T | C | 0.554 | 0.630956 | 0.9893 | *ENOSF1* | 0.261 | 0.024 | 1.69E-27 | 1.62E-21 |
| cg07283778 | TSS200 | rs75588551 | 18 | 725,330 | A | G | 0.122 | 0.901626 | 0.9703 | *ENOSF1* | 0.167 | 0.022 | 1.16E-14 | 7.39E-11 |
| cg15448445 | TSS1500 | rs11113813 | 12 | 108,710,286 | C | G | 0.632 | 0.364196 | 0.9831 | *CMKLR1* | -0.202 | 0.032 | 1.70E-10 | 5.78E-07 |
| cg08110272 | TSS1500 | rs10861891 | 12 | 108,710,323 | C | A | 0.661 | 0.0392288 | 0.9903 | *CMKLR1* | -0.307 | 0.052 | 2.31E-09 | 2.90E-06 |
| cg03612522 | TSS200 | rs4403843 | 12 | 108,707,829 | A | G | 0.662 | 0.0389722 | 0.9685 | *CMKLR1* | -0.102 | 0.018 | 4.62E-09 | 1.14E-05 |
| cg03408433 | TSS1500 | rs11113813 | 12 | 108,710,286 | C | G | 0.632 | 0.364196 | 0.9831 | *CMKLR1* | -0.174 | 0.038 | 2.15E-06 | 1.10E-03 |
| cg25832824 | TSS200 | rs11113813 | 12 | 108,710,286 | C | G | 0.632 | 0.364196 | 0.9831 | *CMKLR1* | -0.076 | 0.017 | 3.74E-06 | 2.70E-03 |
| cg08471037 | TSS200 | rs637718 | 16 | 57,527,946 | A | G | 0.764 | 0.240815 | 0.9642 | *DOK4* | 0.104 | 0.017 | 1.27E-10 | 4.50E-07 |

**Figure 2:**
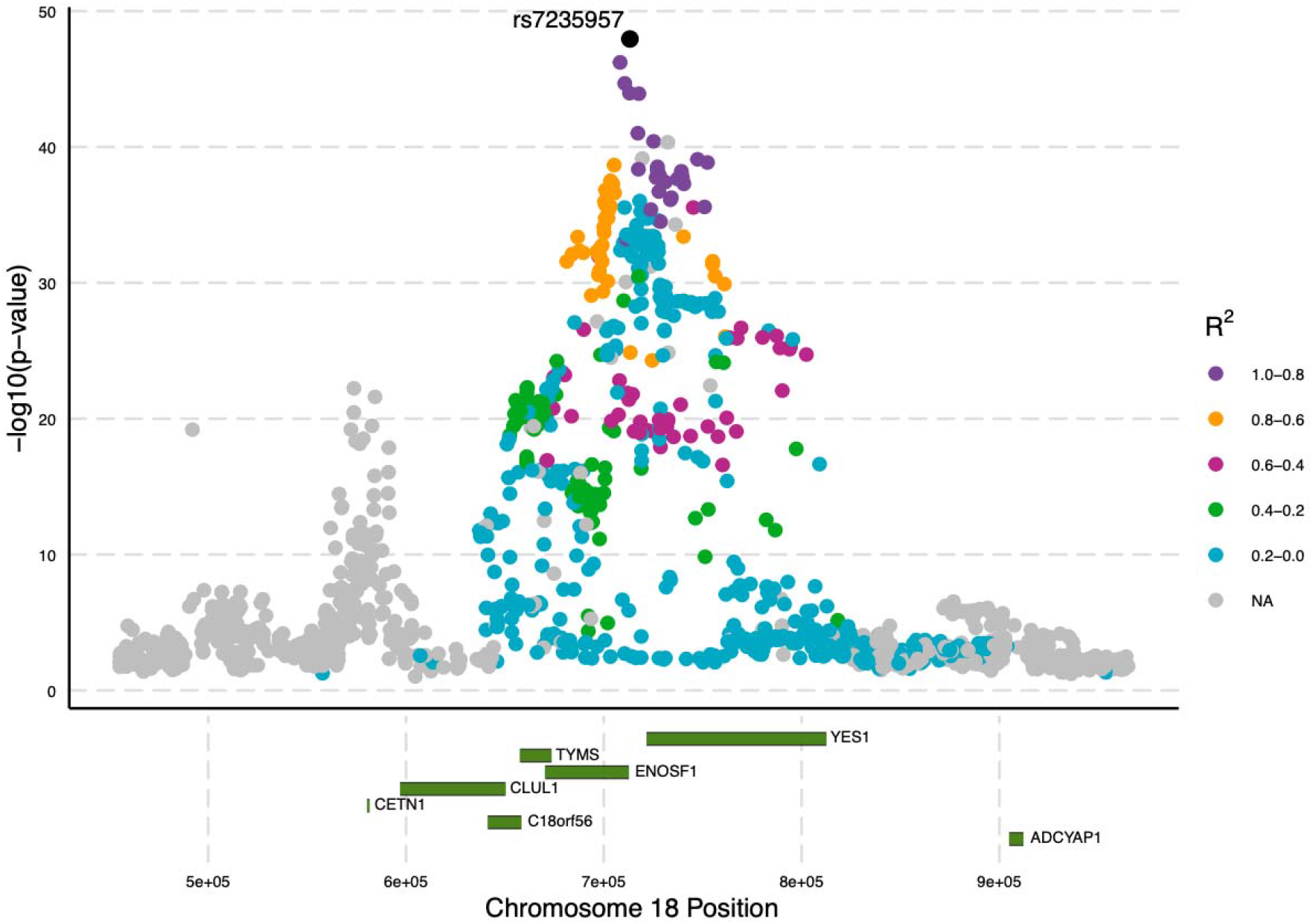
Regional association plot rs7235957 ENOSF1 on chromosome 18. Regional association of variants to DNA methylation in the ENOSF1 promotor region. The strongest association is rs7235957 associated with multiple CpG sites in the ENOSF1 promotor region in carotid artery tissue. Each dot represents a SNP. Lead SNP, highest R^2^, is indicated in black. The X-axis shows the chromosome location relative to 1000 Genomes Project (Nov 2014, Hg19) and refSeq canonical genes (green) from UCSC. The left y-axis shows −log10(p-value) of the association with the CpG site in our region of interest.

### Causal inference of DNAm at 11 shear stress associated genes on cardiovascular risk

Next, we tested the causal effect of differential methylation at 11 shear stress associated genes on cardiovascular risk using our mQTLs (Fig. 3). We used the 9 cis-mQTLs as proxies for the “exposure” DNAm of shear stress associated genes in carotid plaques and we used publicly available GWAS summary statistics of CAD [12] and IS [13] as “outcome” for cardiovascular risk. Overall, CAD analyses show no causal relationship between DNAm of the 11 shear stress associated genes and CAD (inverse variance weighted (IVW): b = −0.007 p = 0.834, Fig. 3a and Table 3). Similarly, IS analyses showed no relationship between DNAm of these genes and IS (wald ratio: b = −0.170 p = 0.317, Table 3). Horizontal pleiotropy was assessed using the MR Egger intercept and showed no pleiotropy (p=0.637). Single SNP analyses of the causal effect of shear stress associated DNAm on CAD also showed no significant results (Fig. 3b, Table 3). Summarizing, causal inference testing of DNAm at the promotor of shear stress associated genes show no significant effect on risk of CAD and IS.

**Figure 3a:**
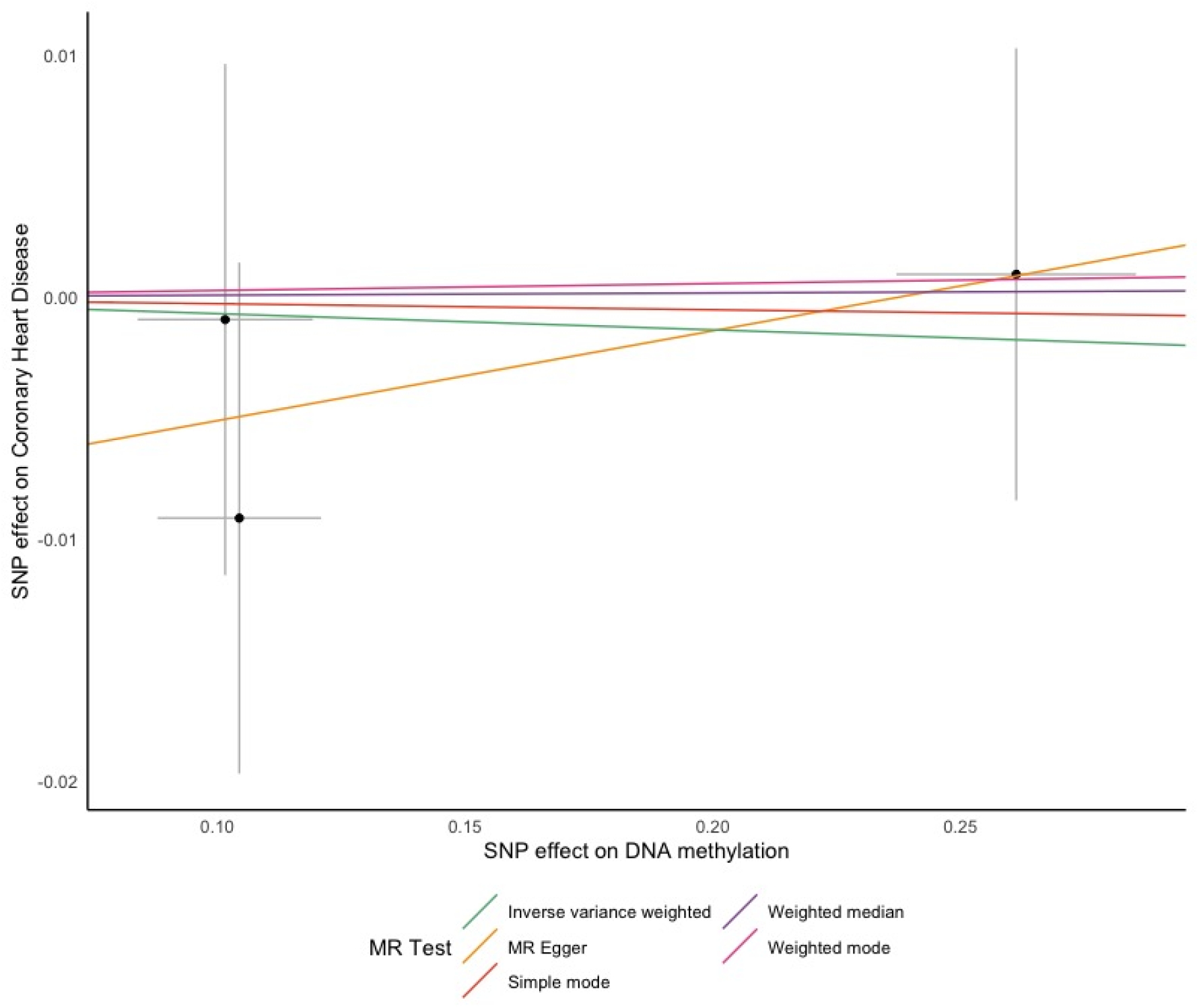
MR Scatterplots of DNA methylation on CAD. (A) 2SMR analysis of 11 shear stress associated genes on cardiovascular disease. We performed 2SMR analysis with plaque mQTLs against the ROIs to test for causality with CAD using GWAS summary-statistics from the CARDIoGRAM-C4D study. Each coloured line corresponds to a performed test indicated by the legend above.

**Figure 3b:**
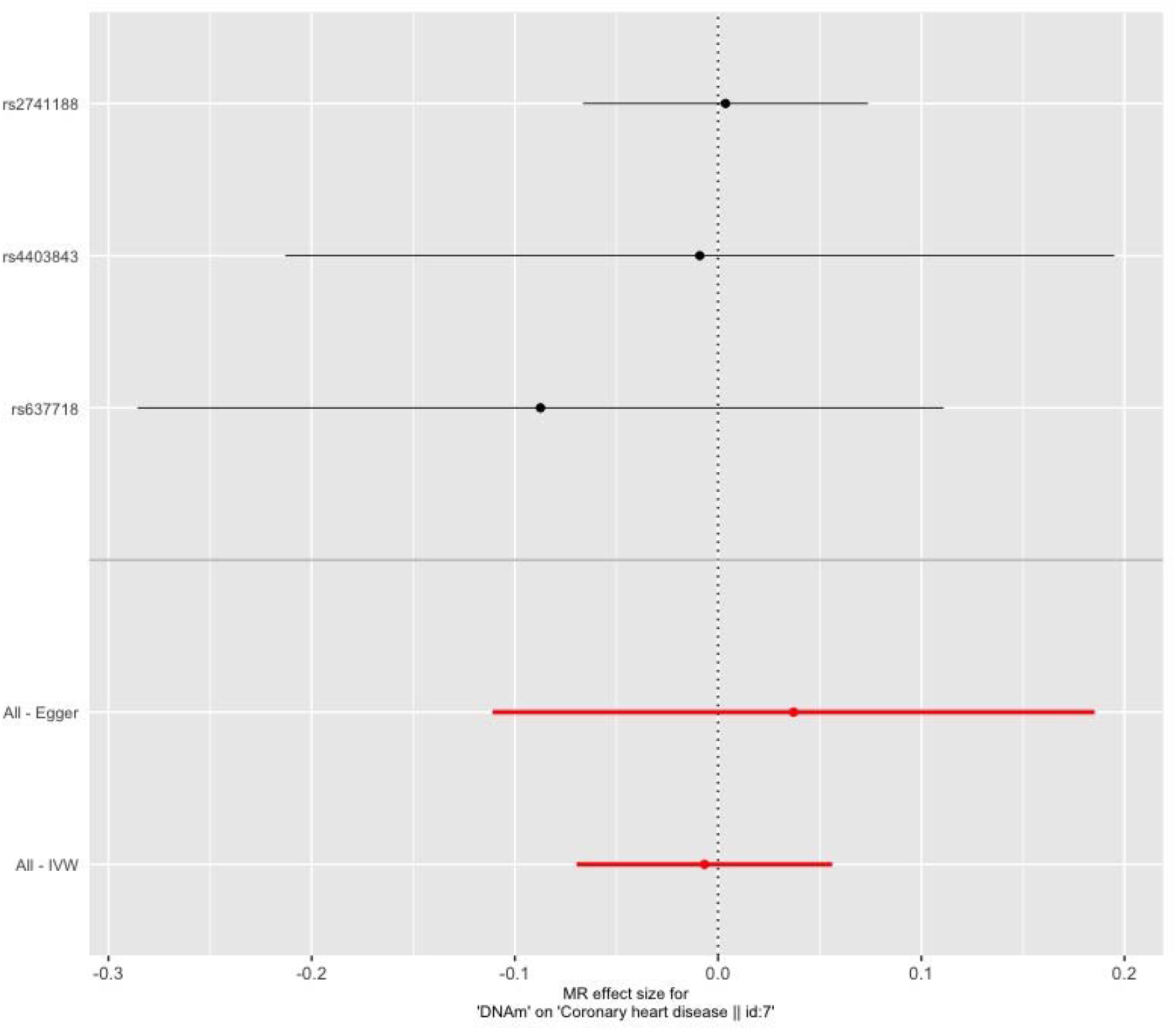
MR Forestplot of DNA methylation on CAD. (B) Single SNP 2SMR analysis of our ROIs mQTLs, as instrumental variants for DNAm of shear stress associated genes on risk of CAD and IS using their respective GWAS summary statistics. Single SNP analysis of shear stress associated DNAm on CAD risk.

**Table 3.** MR results of shear stress associated DNA methylation on CAD and IS. Single SNP and total MR results of shear stress associated DNAm on two cardiovascular outcomes, CAD, using CARDIoGRAM+C4D GWAS summary statistics, and IS, using METASTROKE GWAS summary statistics. Wald Ratio per individual SNP was used for single SNP analyses. (nsnp: number of variants used for MR analysis. SE: standard error of beta).

| Exposure | Outcome | Sample Size | SNP | Beta | SE | P-value |
| --- | --- | --- | --- | --- | --- | --- |
| DNAm | Coronary heart disease | 184,305 | rs2741188 | 0.004 | 0.036 | 0.919 |
|  |  |  | rs4403843 | -0.009 | 0.104 | 0.931 |
|  |  |  | rs637718 | -0.087 | 0.101 | 0.388 |
|  |  |  | All - Inverse variance weighted | -0.007 | 0.032 | 0.834 |
|  |  |  | All - MR Egger | 0.037 | 0.076 | 0.709 |
|  |  |  | *Intercept* | -0.009 |  | 0.637 |
| DNAm | Ischemic stroke | 29,633 | rs4403843 | -0.170 | 0.170 | 0.317 |

## Discussion

We sought to find a causal relationship between differential DNAm of 11 shear stress associated genes in advanced atherosclerotic plaques with cardiovascular disease risk, such as CAD and IS. These genes are associated with initiation of atherosclerosis in mice; here we assessed their role human plaques. We observed no significant overall causal relationship between DNAm of 11 shear stress associated genes in human plaque and increased risk of CAD and IS.

We summit that although methylation of these genes could modulate the initiation of atherosclerosis, collectively it might not result in an increased risk of the ultimate clinical outcome, be it CAD or IS. This could partly be explained by a low sample sizes and lack of replication of the original murine discovery studies, or a suboptimal representation of the human condition by the murine model systems used, i.e. shear stress induced DNAm affects a different set of genes in human compared to mouse models. In addition to these two points, CAD as a proxy for atherosclerosis might not be suitable. CAD is a widespread multifactorial disease rendering the influence of differential DNAm of these 11 shear stress associated genes insignificant. Admittedly, the influence of initial shear stress could be diluted in advanced plaques. Future studies using early stage plaque, from e.g. accidental findings during autopsy, could yield more insight into the role of these 11 genes.

Alternatively, future studies involving endothelial cells, as these are flow-dependent and activation is responsible for atherosclerotic initiation[5,24], could provide more insight in the gene regulatory networks involved in humans and verify the earlier murine results. Such studies could include the design of a shear stress model based on endothelial cells to map of genome-wide differential DNA methylation.

In conclusion, we showed that differential promotor methylation in advanced atherosclerotic plaques of 11 shear stress associated genes, as discovered in mice models, has no significant effect on cardiovascular disease risk. Future research should focus on genome-wide discovery of shear stress associated genes in relevant in vitro models and early stage human plaques.

## Acknowledgements

We acknowledge Lennart Landsmeer, Bas Heijmans, Arjan Boltjes, Michal Mokry, Hester M. den Ruijter, Jessica van Setten, Saskia Haitjema, Gert Jan de Borst, and A. Floriaan Schmidt for fruitful discussions and critical feedback during the study design and writing.

## Author contributions

RM performed research and analysed data.

SWvdL and RM designed the study and wrote the manuscript.

GP provided constructive feedback.

All authors approved the final manuscript

## References

[1] E. Yamamoto, G. Siasos, M. Zaromytidou, A.U. Coskun, L. Xing, K. Bryniarski, T. Zanchin, T. Sugiyama, H. Lee, P.H. Stone, I.-K. Jang, Low Endothelial Shear Stress Predicts Evolution to High-Risk Coronary Plaque Phenotype in the Future: A Serial Optical Coherence Tomography and Computational Fluid Dynamics Study, Circ Cardiovasc Interv. 10 (2017). https://doi.org/10.1161/CIRCINTERVENTIONS.117.005455.

[2] P.H. Stone, A. Maehara, A.U. Coskun, C.C. Maynard, M. Zaromytidou, G. Siasos, I. Andreou, D. Fotiadis, K. Stefanou, M. Papafaklis, L. Michalis, A.J. Lansky, G.S. Mintz, P.W. Serruys, C.L. Feldman, G.W. Stone, Role of Low Endothelial Shear Stress and Plaque Characteristics in the Prediction of Nonculprit Major Adverse Cardiac Events: The PROSPECT Study, JACC Cardiovasc Imaging. 11 (2018) 462–471. https://doi.org/10.1016/j.jcmg.2017.01.031.

[3] J. Dunn, H. Qiu, S. Kim, D. Jjingo, R. Hoffman, C.W. Kim, I. Jang, D.J. Son, D. Kim, C. Pan, Y. Fan, I.K. Jordan, H. Jo, Flow-dependent epigenetic DNA methylation regulates endothelial gene expression and atherosclerosis, J. Clin. Invest. 124 (2014) 3187–3199. https://doi.org/10.1172/JCI74792.

[4] Y. Chan, J.E. Fish, C. D’Abreo, S. Lin, G.B. Robb, A.-M. Teichert, F. Karantzoulis-Fegaras, A. Keightley, B.M. Steer, P.A. Marsden, The cell-specific expression of endothelial nitric-oxide synthase: a role for DNA methylation, J. Biol. Chem. 279 (2004) 35087–35100. https://doi.org/10.1074/jbc.M405063200.

[5] J. Dunn, S. Thabet, H. Jo, Flow-Dependent Epigenetic DNA Methylation in Endothelial Gene Expression and Atherosclerosis, Arterioscler Thromb Vasc Biol. 35 (2015) 1562–1569. https://doi.org/10.1161/ATVBAHA.115.305042.

[6] M.A. Siemelink, S.W. van der Laan, S. Haitjema, I.D. van Koeverden, J. Schaap, M. Wesseling, S.C.A. de Jager, M. Mokry, M. van Iterson, K.F. Dekkers, R. Luijk, H. Foroughi Asl, T. Michoel, J.L.M. Björkegren, E. Aavik, S. Ylä-Herttuala, G.J. de Borst, F.W. Asselbergs, H. El Azzouzi, H.M. den Ruijter, B.T. Heijmans, G. Pasterkamp, Smoking is Associated to DNA Methylation in Atherosclerotic Carotid Lesions, Circ Genom Precis Med. 11 (2018) e002030. https://doi.org/10.1161/CIRCGEN.117.002030.

[7] K.J. Dick, C.P. Nelson, L. Tsaprouni, J.K. Sandling, D. Aïssi, S. Wahl, E. Meduri, P.-E. Morange, F. Gagnon, H. Grallert, M. Waldenberger, A. Peters, J. Erdmann, C. Hengstenberg, F. Cambien, A.H. Goodall, W.H. Ouwehand, H. Schunkert, J.R. Thompson, T.D. Spector, C. Gieger, D.-A. Trégouët, P. Deloukas, N.J. Samani, DNA methylation and body-mass index: a genome-wide analysis, Lancet. 383 (2014) 1990–1998. https://doi.org/10.1016/S0140-6736(13)62674-4.

[8] S. Sayols-Baixeras, I. Subirana, A. Fernández-Sanlés, M. Sentí, C. Lluís-Ganella, J. Marrugat, R. Elosua, DNA methylation and obesity traits: An epigenome-wide association study. The REGICOR study, Epigenetics. 12 (2017) 909–916. https://doi.org/10.1080/15592294.2017.1363951.

[9] X. Xu, S. Su, V.A. Barnes, C. De Miguel, J. Pollock, D. Ownby, H. Shi, H. Zhu, H. Snieder, X. Wang, A genome-wide methylation study on obesity: differential variability and differential methylation, Epigenetics. 8 (2013) 522–533. https://doi.org/10.4161/epi.24506.

[10] C.L. Relton, G. Davey Smith, Two-step epigenetic Mendelian randomization: a strategy for establishing the causal role of epigenetic processes in pathways to disease, Int. J. Epidemiol. 41 (2012) 161–176. https://doi.org/10.1093/ije/dyr233.

[11] G. Davey Smith, G. Hemani, Mendelian randomization: genetic anchors for causal inference in epidemiological studies, Hum. Mol. Genet. 23 (2014) R89–98. https://doi.org/10.1093/hmg/ddu328.

[12] CARDIoGRAMplusC4D Consortium, P. Deloukas, S. Kanoni, C. Willenborg, M. Farrall, T.L. Assimes, J.R. Thompson, E. Ingelsson, D. Saleheen, J. Erdmann, B.A. Goldstein, K. Stirrups, I.R. König, J.-B. Cazier, A. Johansson, A.S. Hall, J.-Y. Lee, C.J. Willer, J.C. Chambers, T. Esko, L. Folkersen, A. Goel, E. Grundberg, A.S. Havulinna, W.K. Ho, J.C. Hopewell, N. Eriksson, M.E. Kleber, K. Kristiansson, P. Lundmark, L.-P. Lyytikäinen, S. Rafelt, D. Shungin, R.J. Strawbridge, G. Thorleifsson, E. Tikkanen, N. Van Zuydam, B.F. Voight, L.L. Waite, W. Zhang, A. Ziegler, D. Absher, D. Altshuler, A.J. Balmforth, I. Barroso, P.S. Braund, C. Burgdorf, S. Claudi-Boehm, D. Cox, M. Dimitriou, R. Do, DIAGRAM Consortium, CARDIOGENICS Consortium, A.S.F. Doney, N. El Mokhtari, P. Eriksson, K. Fischer, P. Fontanillas, A. Franco-Cereceda, B. Gigante, L. Groop, S. Gustafsson, J. Hager, G. Hallmans, B.-G. Han, S.E. Hunt, H.M. Kang, T. Illig, T. Kessler, J.W. Knowles, G. Kolovou, J. Kuusisto, C. Langenberg, C. Langford, K. Leander, M.-L. Lokki, A. Lundmark, M.I. McCarthy, C. Meisinger, O. Melander, E. Mihailov, S. Maouche, A.D. Morris, M. Müller-Nurasyid, MuTHER Consortium, K. Nikus, J.F. Peden, N.W. Rayner, A. Rasheed, S. Rosinger, D. Rubin, M.P. Rumpf, A. Schäfer, M. Sivananthan, C. Song, A.F.R. Stewart, S.-T. Tan, G. Thorgeirsson, C.E. van der Schoot, P.J. Wagner, Wellcome Trust Case Control Consortium, G.A. Wells, P.S. Wild, T.-P. Yang, P. Amouyel, D. Arveiler, H. Basart, M. Boehnke, E. Boerwinkle, P. Brambilla, F. Cambien, A.L. Cupples, U. de Faire, A. Dehghan, P. Diemert, S.E. Epstein, A. Evans, M.M. Ferrario, J. Ferrières, D. Gauguier, A.S. Go, A.H. Goodall, V. Gudnason, S.L. Hazen, H. Holm, C. Iribarren, Y. Jang, M. Kähönen, F. Kee, H.-S. Kim, N. Klopp, W. Koenig, W. Kratzer, K. Kuulasmaa, M. Laakso, R. Laaksonen, J.-Y. Lee, L. Lind, W.H. Ouwehand, S. Parish, J.E. Park, N.L. Pedersen, A. Peters, T. Quertermous, D.J. Rader, V. Salomaa, E. Schadt, S.H. Shah, J. Sinisalo, K. Stark, K. Stefansson, D.-A. Trégouët, J. Virtamo, L. Wallentin, N. Wareham, M.E. Zimmermann, M.S. Nieminen, C. Hengstenberg, M.S. Sandhu, T. Pastinen, A.-C. Syvänen, G.K. Hovingh, G. Dedoussis, P.W. Franks, T. Lehtimäki, A. Metspalu, P.A. Zalloua, A. Siegbahn, S. Schreiber, S. Ripatti, S.S. Blankenberg, M. Perola, R. Clarke, B.O. Boehm, C. O’Donnell, M.P. Reilly, W. März, R. Collins, S. Kathiresan, A. Hamsten, J.S. Kooner, U. Thorsteinsdottir, J. Danesh, C.N.A. Palmer, R. Roberts, H. Watkins, H. Schunkert, N.J. Samani, Large-scale association analysis identifies new risk loci for coronary artery disease, Nat. Genet. 45 (2013) 25–33. https://doi.org/10.1038/ng.2480.

[13] R. Malik, G. Chauhan, M. Traylor, M. Sargurupremraj, Y. Okada, A. Mishra, L. Rutten-Jacobs, A.-K. Giese, S.W. van der Laan, S. Gretarsdottir, C.D. Anderson, M. Chong, H.H.H. Adams, T. Ago, P. Almgren, P. Amouyel, H. Ay, T.M. Bartz, O.R. Benavente, S. Bevan, G.B. Boncoraglio, R.D. Brown, A.S. Butterworth, C. Carrera, C.L. Carty, D.I. Chasman, W.-M. Chen, J.W. Cole, A. Correa, I. Cotlarciuc, C. Cruchaga, J. Danesh, P.I.W. de Bakker, A.L. DeStefano, M. den Hoed, Q. Duan, S.T. Engelter, G.J. Falcone, R.F. Gottesman, R.P. Grewal, V. Gudnason, S. Gustafsson, J. Haessler, T.B. Harris, A. Hassan, A.S. Havulinna, S.R. Heckbert, E.G. Holliday, G. Howard, F.-C. Hsu, H.I. Hyacinth, M.A. Ikram, E. Ingelsson, M.R. Irvin, X. Jian, J. Jiménez-Conde, J.A. Johnson, J.W. Jukema, M. Kanai, K.L. Keene, B.M. Kissela, D.O. Kleindorfer, C. Kooperberg, M. Kubo, L.A. Lange, C.D. Langefeld, C. Langenberg, L.J. Launer, J.-M. Lee, R. Lemmens, D. Leys, C.M. Lewis, W.-Y. Lin, A.G. Lindgren, E. Lorentzen, P.K. Magnusson, J. Maguire, A. Manichaikul, P.F. McArdle, J.F. Meschia, B.D. Mitchell, T.H. Mosley, M.A. Nalls, T. Ninomiya, M.J. O’Donnell, B.M. Psaty, S.L. Pulit, K. Rannikmäe, A.P. Reiner, K.M. Rexrode, K. Rice, S.S. Rich, P.M. Ridker, N.S. Rost, P.M. Rothwell, J.I. Rotter, T. Rundek, R.L. Sacco, S. Sakaue, M.M. Sale, V. Salomaa, B.R. Sapkota, R. Schmidt, C.O. Schmidt, U. Schminke, P. Sharma, A. Slowik, C.L.M. Sudlow, C. Tanislav, T. Tatlisumak, K.D. Taylor, V.N.S. Thijs, G. Thorleifsson, U. Thorsteinsdottir, S. Tiedt, S. Trompet, C. Tzourio, C.M. van Duijn, M. Walters, N.J. Wareham, S. Wassertheil-Smoller, J.G. Wilson, K.L. Wiggins, Q. Yang, S. Yusuf, J.C. Bis, T. Pastinen, A. Ruusalepp, E.E. Schadt, S. Koplev, J.L.M. Björkegren, V. Codoni, M. Civelek, N.L. Smith, D.A. Trégouët, I.E. Christophersen, C. Roselli, S.A. Lubitz, P.T. Ellinor, E.S. Tai, J.S. Kooner, N. Kato, J. He, P. van der Harst, P. Elliott, J.C. Chambers, F. Takeuchi, A.D. Johnson, D.K. Sanghera, O. Melander, C. Jern, D. Strbian, I. Fernandez-Cadenas, W.T. Longstreth, A. Rolfs, J. Hata, D. Woo, J. Rosand, G. Pare, J.C. Hopewell, D. Saleheen, K. Stefansson, B.B. Worrall, S.J. Kittner, S. Seshadri, M. Fornage, H.S. Markus, J.M.M. Howson, Y. Kamatani, S. Debette, M. Dichgans, R. Malik, G. Chauhan, M. Traylor, M. Sargurupremraj, Y. Okada, A. Mishra, L. Rutten-Jacobs, A.-K. Giese, S.W. van der Laan, S. Gretarsdottir, C.D. Anderson, M. Chong, H.H.H. Adams, T. Ago, P. Almgren, P. Amouyel, H. Ay, T.M. Bartz, O.R. Benavente, S. Bevan, G.B. Boncoraglio, R.D. Brown, A.S. Butterworth, C. Carrera, C.L. Carty, D.I. Chasman, W.-M. Chen, J.W. Cole, A. Correa, I. Cotlarciuc, C. Cruchaga, J. Danesh, P.I.W. de Bakker, A.L. DeStefano, M. den Hoed, Q. Duan, S.T. Engelter, G.J. Falcone, R.F. Gottesman, R.P. Grewal, V. Gudnason, S. Gustafsson, J. Haessler, T.B. Harris, A. Hassan, A.S. Havulinna, S.R. Heckbert, E.G. Holliday, G. Howard, F.-C. Hsu, H.I. Hyacinth, M.A. Ikram, E. Ingelsson, M.R. Irvin, X. Jian, J. Jiménez-Conde, J.A. Johnson, J.W. Jukema, M. Kanai, K.L. Keene, B.M. Kissela, D.O. Kleindorfer, C. Kooperberg, M. Kubo, L.A. Lange, C.D. Langefeld, C. Langenberg, L.J. Launer, J.-M. Lee, R. Lemmens, D. Leys, C.M. Lewis, W.-Y. Lin, A.G. Lindgren, E. Lorentzen, P.K. Magnusson, J. Maguire, A. Manichaikul, P.F. McArdle, J.F. Meschia, B.D. Mitchell, T.H. Mosley, M.A. Nalls, T. Ninomiya, M.J. O’Donnell, B.M. Psaty, S.L. Pulit, K. Rannikmäe, A.P. Reiner, K.M. Rexrode, K. Rice, S.S. Rich, P.M. Ridker, N.S. Rost, P.M. Rothwell, J.I. Rotter, T. Rundek, R.L. Sacco, S. Sakaue, M.M. Sale, V. Salomaa, B.R. Sapkota, R. Schmidt, C.O. Schmidt, U. Schminke, P. Sharma, A. Slowik, C.L.M. Sudlow, C. Tanislav, T. Tatlisumak, K.D. Taylor, V.N.S. Thijs, G. Thorleifsson, U. Thorsteinsdottir, S. Tiedt, S. Trompet, C. Tzourio, C.M. van Duijn, M. Walters, N.J. Wareham, S. Wassertheil-Smoller, J.G. Wilson, K.L. Wiggins, Q. Yang, S. Yusuf, N. Amin, H.S. Aparicio, D.K. Arnett, J. Attia, A.S. Beiser, C. Berr, J.E. Buring, M. Bustamante, V. Caso, Y.-C. Cheng, S.H. Choi, A. Chowhan, N. Cullell, J.-F. Dartigues, H. Delavaran, P. Delgado, M. Dörr, G. Engström, I. Ford, W.S. Gurpreet, A. Hamsten, L. Heitsch, A. Hozawa, L. Ibanez, A. Ilinca, M. Ingelsson, M. Iwasaki, R.D. Jackson, K. Jood, P. Jousilahti, S. Kaffashian, L. Kalra, M. Kamouchi, T. Kitazono, O. Kjartansson, M. Kloss, P.J. Koudstaal, J. Krupinski, D.L. Labovitz, C.C. Laurie, C.R. Levi, L. Li, L. Lind, C.M. Lindgren, V. Lioutas, Y.M. Liu, O.L. Lopez, H. Makoto, N. Martinez-Majander, K. Matsuda, N. Minegishi, J. Montaner, A.P. Morris, E. Muiño, M. Müller-Nurasyid, B. Norrving, S. Ogishima, E.A. Parati, L.R. Peddareddygari, N.L. Pedersen, J. Pera, M. Perola, A. Pezzini, S. Pileggi, R. Rabionet, I. Riba-Llena, M. Ribasés, J.R. Romero, J. Roquer, A.G. Rudd, A.-P. Sarin, R. Sarju, C. Sarnowski, M. Sasaki, C.L. Satizabal, M. Satoh, N. Sattar, N. Sawada, G. Sibolt, Á. Sigurdsson, A. Smith, K. Sobue, C. Soriano-Tárraga, T. Stanne, O.C. Stine, D.J. Stott, K. Strauch, T. Takai, H. Tanaka, K. Tanno, A. Teumer, L. Tomppo, N.P. Torres-Aguila, E. Touze, S. Tsugane, A.G. Uitterlinden, E.M. Valdimarsson, S.J. van der Lee, H. Völzke, K. Wakai, D. Weir, S.R. Williams, C.D.A. Wolfe, Q. Wong, H. Xu, T. Yamaji, D.K. Sanghera, O. Melander, C. Jern, D. Strbian, I. Fernandez-Cadenas, W.T. Longstreth, A. Rolfs, J. Hata, D. Woo, J. Rosand, G. Pare, J.C. Hopewell, D. Saleheen, K. Stefansson, B.B. Worrall, S.J. Kittner, S. Seshadri, M. Fornage, H.S. Markus, J.M.M. Howson, Y. Kamatani, S. Debette, M. Dichgans, AFGen Consortium, Cohorts for Heart and Aging Research in Genomic Epidemiology (CHARGE) Consortium, International Genomics of Blood Pressure (iGEN-BP) Consortium, INVENT Consortium, STARNET, BioBank Japan Cooperative Hospital Group, COMPASS Consortium, EPIC-CVD Consortium, EPIC-InterAct Consortium, International Stroke Genetics Consortium (ISGC), METASTROKE Consortium, Neurology Working Group of the CHARGE Consortium, NINDS Stroke Genetics Network (SiGN), UK Young Lacunar DNA Study, MEGASTROKE Consortium, MEGASTROKE Consortium:, Multiancestry genome-wide association study of 520,000 subjects identifies 32 loci associated with stroke and stroke subtypes, Nat. Genet. 50 (2018) 524–537. https://doi.org/10.1038/s41588-018-0058-3.

[14] E. Aavik, M. Babu, S. Ylä-Herttuala, DNA methylation processes in atheosclerotic plaque, Atherosclerosis. 281 (2019) 168–179. https://doi.org/10.1016/j.atherosclerosis.2018.12.006.

[15] B.A.N. Verhoeven, E. Velema, A.H. Schoneveld, J.P.P.M. de Vries, P. de Bruin, C.A. Seldenrijk, D.P.V. de Kleijn, E. Busser, Y. van der Graaf, F. Moll, G. Pasterkamp, Athero-express: differential atherosclerotic plaque expression of mRNA and protein in relation to cardiovascular events and patient characteristics. Rationale and design, Eur. J. Epidemiol. 19 (2004) 1127–1133.

[16] S.W. van der Laan, H. Foroughi Asl, P. van den Borne, J. van Setten, M.E.M. van der Perk, S.M. van de Weg, A.H. Schoneveld, D.P.V. de Kleijn, T. Michoel, J.L.M. Björkegren, H.M. den Ruijter, F.W. Asselbergs, P.I.W. de Bakker, G. Pasterkamp, Variants in ALOX5, ALOX5AP and LTA4H are not associated with atherosclerotic plaque phenotypes: the Athero-Express Genomics Study, Atherosclerosis. 239 (2015) 528–538. https://doi.org/10.1016/j.atherosclerosis.2015.01.018.

[17] C.C. Laurie, K.F. Doheny, D.B. Mirel, E.W. Pugh, L.J. Bierut, T. Bhangale, F. Boehm, N.E. Caporaso, M.C. Cornelis, H.J. Edenberg, S.B. Gabriel, E.L. Harris, F.B. Hu, K.B. Jacobs, P. Kraft, M.T. Landi, T. Lumley, T.A. Manolio, C. McHugh, I. Painter, J. Paschall, J.P. Rice, K.M. Rice, X. Zheng, B.S. Weir, GENEVA Investigators, Quality control and quality assurance in genotypic data for genome-wide association studies, Genet. Epidemiol. 34 (2010) 591–602. https://doi.org/10.1002/gepi.20516.

[18] 1000 Genomes Project Consortium, G.R. Abecasis, D. Altshuler, A. Auton, L.D. Brooks, R.M. Durbin, R.A. Gibbs, M.E. Hurles, G.A. McVean, A map of human genome variation from population-scale sequencing, Nature. 467 (2010) 1061–1073. https://doi.org/10.1038/nature09534.

[19] D.I. Boomsma, C. Wijmenga, E.P. Slagboom, M.A. Swertz, L.C. Karssen, A. Abdellaoui, K. Ye, V. Guryev, M. Vermaat, F. van Dijk, L.C. Francioli, J.J. Hottenga, J.F.J. Laros, Q. Li, Y. Li, H. Cao, R. Chen, Y. Du, N. Li, S. Cao, J. van Setten, A. Menelaou, S.L. Pulit, J.Y. Hehir-Kwa, M. Beekman, C.C. Elbers, H. Byelas, A.J.M. de Craen, P. Deelen, M. Dijkstra, J.T. den Dunnen, P. de Knijff, J. Houwing-Duistermaat, V. Koval, K. Estrada, A. Hofman, A. Kanterakis, D. van Enckevort, H. Mai, M. Kattenberg, E.M. van Leeuwen, P.B.T. Neerincx, B. Oostra, F. Rivadeneira, E.H.D. Suchiman, A.G. Uitterlinden, G. Willemsen, B.H. Wolffenbuttel, J. Wang, P.I.W. de Bakker, G.-J. van Ommen, C.M. van Duijn, The Genome of the Netherlands: design, and project goals, Eur. J. Hum. Genet. 22 (2014) 221–227. https://doi.org/10.1038/ejhg.2013.118.

[20] B.N. Howie, P. Donnelly, J. Marchini, A flexible and accurate genotype imputation method for the next generation of genome-wide association studies, PLoS Genet. 5 (2009) e1000529. https://doi.org/10.1371/journal.pgen.1000529.

[21] Jaccoschaap, S.W.V.D. Laan, Swvanderlaan/Qtltoolkit: T’Pol, Zenodo, 2017. https://doi.org/10.5281/ZENODO.1040185.

[22] O. Delaneau, H. Ongen, A.A. Brown, A. Fort, N.I. Panousis, E.T. Dermitzakis, A complete tool set for molecular QTL discovery and analysis, Nat Commun. 8 (2017) 15452. https://doi.org/10.1038/ncomms15452.

[23] J. Bowden, G. Davey Smith, S. Burgess, Mendelian randomization with invalid instruments: effect estimation and bias detection through Egger regression, Int J Epidemiol. 44 (2015) 512–525. https://doi.org/10.1093/ije/dyv080.

[24] P.N. Hopkins, Molecular biology of atherosclerosis, Physiol. Rev. 93 (2013) 1317–1542. https://doi.org/10.1152/physrev.00004.2012.

