## Supplemental Material for "Exploring the Causal Effects of Shear Stress Associated DNA Methylation on Cardiovascular Risk"

**Appendix A: Supplementary data**

**Supplemental table 1: mQTL regions of interest (ROIs)**

|  | | | **Chromosome Locations** | | | | | | | |
| --- | --- | --- | --- | --- | --- | --- | --- | --- | --- | --- |
| **Gene** | **chr** | **Strand** | **TSS** | **2000TSS** | **exon** | **ROI Start** | **ROI Stop** | **QTL start** | **QTL stop** | **QTL region size** |
| HOXA5 | chr7 | - | 27,183,287 | 27,185,287 | 27,182,665 | 27,182,665 | 27,185,287 | 26,932,665 | 27,435,287 | 502,622 |
| TMEM184B | chr16 | - | 38,669,040 | 38,671,040 | 38,668,890 | 38,668,890 | 38,671,040 | 38,418,890 | 38,921,040 | 502,150 |
| ADAMTSL5 | chr19 | - | 1,513,188 | 1,515,188 | 1,512,962 | 1,512,962 | 1,515,188 | 1,262,962 | 1,765,188 | 502,226 |
| KLF4 | chr9 | - | 110,251,927 | 110,253,927 | 110,251,449 | 110,251,449 | 110,253,927 | 110,001,449 | 110,503,927 | 502,478 |
| KLF3 | chr4 | + | 38,665,817 | 38,663,817 | 38,666,082 | 38,663,817 | 38,666,082 | 38,413,817 | 38,916,082 | 502,265 |
| CMKLR1 | chr12 | - | 108,733,094 | 108,735,094 | 108,732,804 | 108,732,804 | 108,735,094 | 108,482,804 | 108,985,094 | 502,290 |
| PKP4 | chr2 | + | 159,313,611 | 159,311,611 | 159,313,730 | 159,311,611 | 159,313,730 | 159,061,611 | 159,563,730 | 502,119 |
| ACVRL1 | chr12 | + | 52,301,202 | 52,298,202 | 52,301,479 | 52,298,202 | 52,301,479 | 52,048,202 | 52,551,479 | 503,277 |
| DOK4 | chr16 | - | 57,520,407 | 57,522,407 | 57,520,217 | 57,520,217 | 57,522,407 | 57,270,217 | 57,772,407 | 502,190 |
| SPRY2 | chr13 | - | 80,915,086 | 80,917,086 | 80,914,757 | 80,914,757 | 80,917,086 | 80,664,757 | 81,167,086 | 502,329 |
| ENOSF1 | chr18 | - | 712,676 | 715,676 | 712,504 | 712,504 | 715,676 | 462,504 | 965,676 | 503,172 |

The regions used for mQTL discovery per shear stress associated gene. A region is established by a 250 kb region around the outermost CpGs of the -2000 TSS to first exon of each gene. Gene; refSeq canonical genes from UCSC, Chr; Chromosome, Str; forward (+) or reverse (-) strand, TSS; transcription start site, 2000TSS; -2000 transcription start site, ROI start/stop; region of interest (promotor region), QTL start/stop; region used for mQTL analysis, Chromosome Locations; chromosome locations relative to 1000 Genomes Project (Nov 2014, Hg19).

**Supplemental excel table**

Nominal and permutation mQTL results

**Supplemental figure 1: mQTL power estimation**

mQTL power estimation for our analysis. We performed 1,674,865 SNP tests (nSNP) over a total of 1,674,865 CpGs (before filtering for CpGs within our ROI) for 442 samples. A minor allele frequency (MAF) > 0.06 has a proper power estimation of 85% and higher.

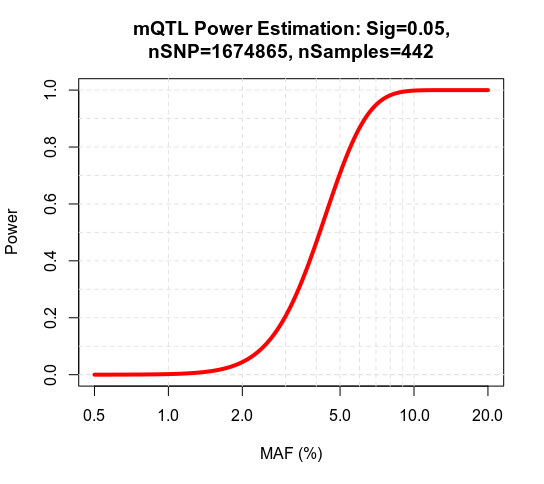
